# Hierarchical Frequency Tagging reveals neural markers of predictive coding under varying uncertainty

**DOI:** 10.1101/081349

**Authors:** Noam Gordon, Roger Koenig-Robert, Naotsugu Tsuchiya, Jeroen van Boxtel, Jakob Hohwy

## Abstract

Understanding the integration of top-down and bottom-up signals is essential for the study of perception. Current accounts of predictive coding describe this in terms of interactions between state units encoding expectations or predictions, and error units encoding prediction error. However, direct neural evidence for such interactions has not been well established. To achieve this, we combined EEG methods that preferentially tag different levels in the visual hierarchy: Steady State Visual Evoked Potential (SSVEP at 10Hz, tracking bottom-up signals) and Semantic Wavelet-Induced Frequency Tagging (SWIFT at 1.3Hz tracking top-down signals). Importantly, we examined intermodulation components (IM, e.g., 11.3Hz) as a measure of integration between these signals. To examine the influence of expectation and predictions on the nature of such integration, we constructed 50-second movie streams and modulated expectation levels for upcoming stimuli by varying the proportion of images presented across trials. We found SWIFT, SSVEP and IM signals to differ in important ways. SSVEP was strongest over occipital electrodes and was not modified by certainty. Conversely, SWIFT signals were evident over temporo- and parieto-occipital areas and decreased as a function of increasing certainty levels. Finally, IMs were evident over occipital electrodes and increased as a function of certainty. These results link SSVEP, SWIFT and IM signals to sensory evidence, predictions, prediction errors and hypothesis-testing - the core elements of predictive coding. These findings provide neural evidence for the integration of top-down and bottom-up information in perception, opening new avenues to studying such interactions in perception while constraining neuronal models of predictive coding.

**SIGNIFICANCE STATEMENT:** There is a growing understanding that both top-down and bottom-up signals underlie perception. But how do these signals interact? And how does this process depend on the signals’ probabilistic properties? ‘Predictive coding’ theories of perception describe this in terms how well top-down predictions fit with bottom-up sensory input. Identifying neural markers for such signal integration is therefore essential for the study of perception and predictive coding theories in particular. The novel Hierarchical Frequency Tagging method simultaneously tags top-down and bottom-up signals in EEG recordings, while obtaining a measure for the level of integration between these signals. Our results suggest that top-down predictions indeed integrate with bottom-up signals in a manner that is modulated by the predictability of the sensory input.

## 1. INTRODUCTION

Perception is increasingly being understood to arise by means of cortical integration of ‘bottom-up’ or sensory-driven signals and ‘top-down’ information. Prior experience, expectations and knowledge about the world allow for the formation of priors or hypotheses about the state of the external world (i.e., the causes of the sensory input) that help resolve sensory ambiguity. Such neuronal representations, or ‘state-units’ can then be tested and optimised in light of new sensory evidence. Contemporary accounts treat these ideas in terms of Bayesian inference and predictive coding (1–4).

That perception is essentially an inferential process is supported by many behavioural findings demonstrating the significant role of contextual information (5–8) and of top-down signals (9–12) in perception. Several studies additionally suggest different neural measures of feedforward and feedback signals (13) primarily in terms of their characteristic oscillatory frequency bands (14–19).

However, studying the neural basis of perception requires not only distinguishing between top-down and bottom-up signals but also examining the actual integration between such signals.

This is particularly important for predictive coding, which hypothesizes such integration as a mechanism for prediction error minimization that is marked by the probabilistic properties of predictions and prediction errors. Hence, the goals of this study were to simultaneously tag top-down and bottom-up signals and then identify a direct neural marker for the integration of these signals during visual perception and, further, to examine if, and how, such a marker is modulated by the strength of prior expectations.

In order to differentiate between top-down signals related to predictions, bottom-up signals related to the accumulation of sensory evidence, and the interaction between such signals, we developed the Hierarchical Frequency Tagging (HFT) paradigm in which two frequency tagging methods are combined in the visual domain in a hierarchical manner. To preferentially track top-down signals (i.e., putative prediction signals) we used semantic wavelet induced frequency tagging (SWIFT) that has been shown to constantly activate low-level visual areas while periodically engaging high-level visual areas (thus, selectively tagging the high-level visual areas; (20, 21)). To simultaneously track bottom-up signals (i.e., sensory evidence) we used classic frequency tagging, or so called steady state visual evoked potentials (SSVEP) (22, 23). We combined the two methods by presenting SWIFT-modulated images at 1.3HZ while modulating the global luminance of the stimulus at 10Hz to elicit SSVEP (See Methods for details). Critically, we hypothesized that intermodulation (IM) components would appear as a marker of integration between these differentially tagged signals.

Intermodulation is a common phenomenon manifesting in non-linear systems. When the input signal is comprised of more than one fundamental frequency (e.g., *F1* and *F2*) that interact within a non-linear system, the response output will show additional frequencies as linear combinations of the input frequencies (e.g., *f1* + *f2*, *f1* - *f2*, etc.) (note that throughout the paper we denote stimulus frequencies with capital letters (e.g., F1) and response frequencies with small letters (e.g., f1)). Intermodulation components in EEG recordings have been used to study non-linear interactions in the visual system (24–26), with some recent applications for the study of high-level visual-object recognition systems (27–29). However, instead of tagging two ‘bottom-up’ signals, our paradigm was designed to enable the examination of the integration between both bottom-up *and* top-down inputs to the lower visual areas. We hypothesized that the hierarchical combination of SSVEP and SWIFT (HFT) will allow us to track sensory-driven signals as SSVEP and prediction-driven signals as SWIFT, while having the integration between these signals as the IM components. We therefore additionally hypothesised that the IM components will be modulated by the probabilistic properties (e.g., the certainty or predictability) of the stimuli giving rise to the SSVEP and SWIFT signals.

To test these hypotheses and examine whether the IMs observed in our paradigm indeed reflect the hierarchical integration of predictions and sensory evidence in perceptual inference, we manipulated levels of expectation across trials. Expectation reflects the continuous process of probabilistic learning about what is possible or probable in the forthcoming sensory environment (30) and therefore plays a central role in predictive coding. Indeed, various studies have demonstrated the relationship between stimulus predictability and neural responses (10, 31). Accordingly, we varied the levels of expectation (i.e. certainty regarding upcoming stimuli) across trials in order to modulate prediction-driven signals (see Methods for further details). In this way, we aimed not only to find neural markers for the integration of sensory-driven and prediction-driven signals, but also to examine how the process is modulated by expectation and uncertainty - core elements in the predictive coding framework.

## 2. RESULTS

Participants were presented with 50-sec ‘movie’ streams in which either a house or a face image appeared briefly at a frequency of 1.3Hz. Each 50-sec trial was constructed using one face and one house image randomly selected from a pool of images. Images were scrambled using two frequency tagging methods - SWIFT and SSVEP - that differentially tag areas in the cortical hierarchy (Fig. 1). Prior to each trial, participants were instructed to count the number of times one of the two images appeared in the trial (either the house or the face image) and they reported their response at the end of each trial. The proportion of images changed over trials, ranging from trials in which both images appeared in nearly half the cycles (referred to as ‘low certainty’ trials) to trials in which one of the images appeared in nearly all cycles (referred to as ‘high certainty’ trials).

**Figure 1.**
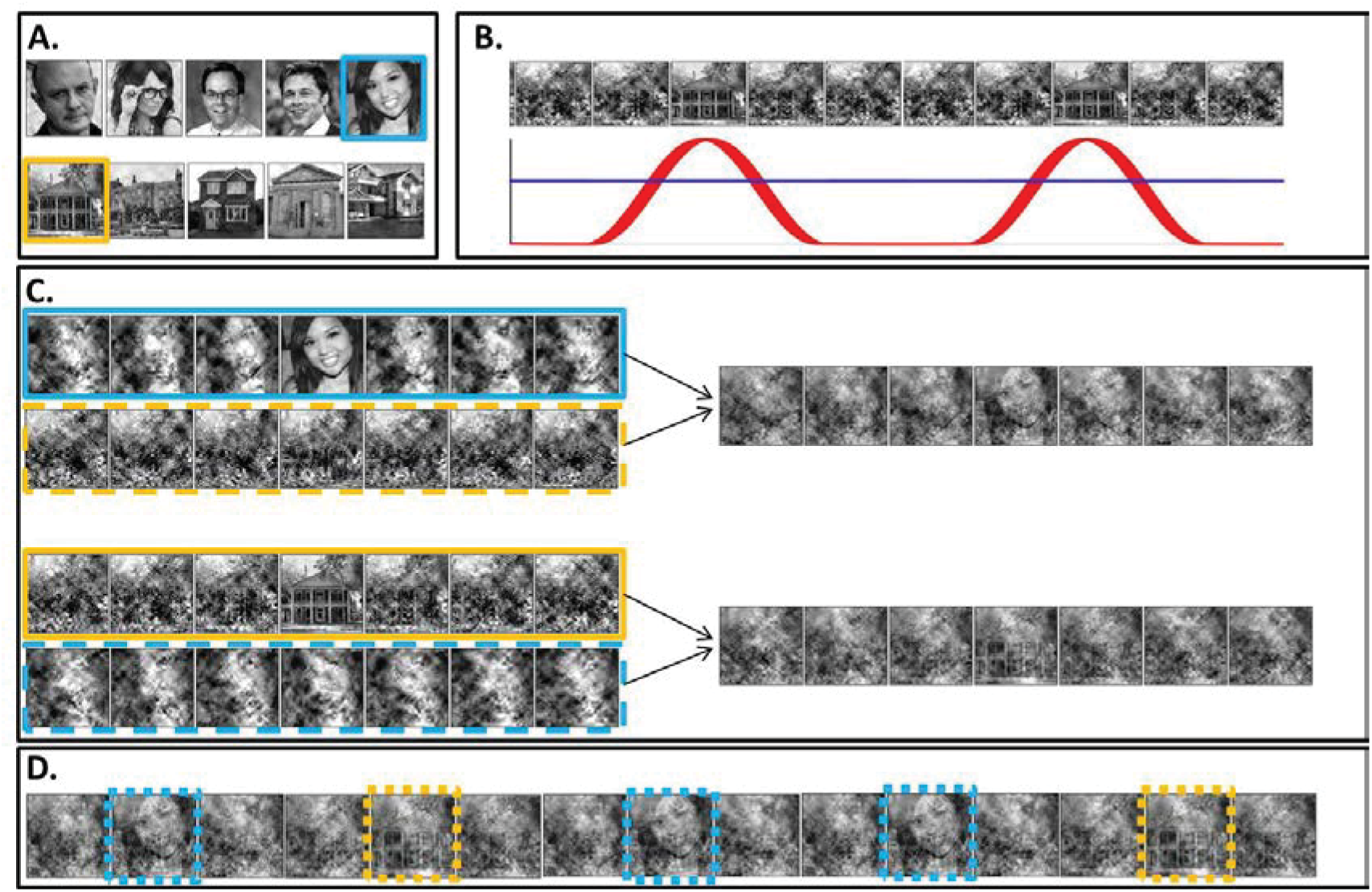
Stimuli construction. Schematic illustration of stimuli construction. (A) A pool of 28 face and 28 house images were used in the paradigm. (B) The SWIFT principle. Cyclic local-contour scrambling in the wavelet-domain allows us to modulate the semantics of the image at a given frequency (i.e. the tagging-frequency, F2=1.3hz, illustrated by the solid red line) while keeping low-level principal physical attributes constant over time (illustrated by the dashed blue line) (C) Each trial (50 seconds) was constructed using one SWIFT cycle (~769 ms) of a randomly chosen face image (blue solid rectangle) and one SWIFT cycle of a randomly chosen house image (orange solid rectangle). For each SWIFT cycle, a corresponding ‘noise’ SWIFT cycle was created based on one of the scrambled frames of the original SWIFT cycle (orange and blue dashed rectangles). Superimposition of the original (solid rectangles) and noise (dashed rectangles) SWIFT cycles ensures similar principal local physical properties across all SWIFT frames. (D) The two SWIFT cycles (house and face) were presented repeatedly in a pseudo-random order for a total of 65 cycles. The resulting trial was a 50 second movie in which images peaked in a cyclic manner (F2=1.3Hz). Finally, a global sinusoidal contrast modulation at F1=10Hz was applied on to the whole movie to evoke the SSVEP.

Having assured that participants were able to perform the task (Fig. 6), we first verified whether our two frequency-tagging methods were indeed able to entrain brain activity, and whether we could observe intermodulation (IM) components. Figure 2 shows the results of the Fourier transform (FFT) averaged across all 64 electrodes, trials and participants (N=17). Importantly, significant peaks can be seen at both tagging frequencies (f1=10Hz and f2=1.3Hz) and their harmonics (n1f1 and n2f2; red and pink solid lines in Figure 2) and at various IM components (n1f1+n2f2; orange dashed lines in Figure 2) (one sample *t*-test, FDR-adjusted p < 0.01 for frequencies of interest in the range of 1Hz-40Hz).

**Figure 2.**
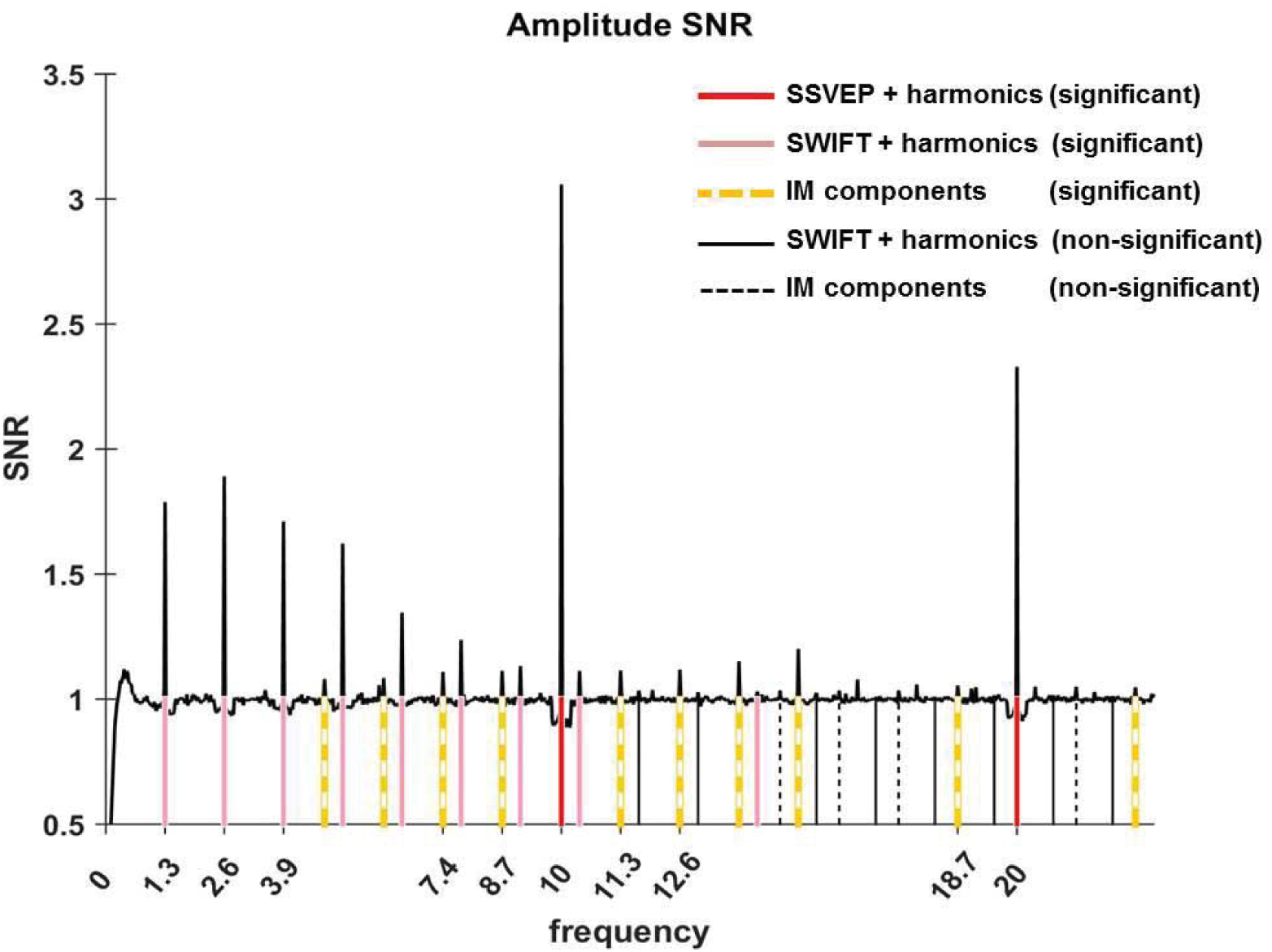
Amplitude SNR spectra. Amplitude SNRs (see Methods for the definition of SNR), averaged across all electrodes, trials and participants, are shown for frequencies up to 23Hz. Peaks can be seen at the tagging frequencies, their harmonics and at IM components. Solid red lines mark the SSVEP frequency and its harmonic (10Hz and 20Hz, both with SNRs significantly greater than one). Solid pink lines mark the SWIFT frequency and harmonics with SNRs significantly greater than one (n2f2 where n2=1,2,3…8 and 11). Solid black lines mark SWIFT harmonics with SNRs not significantly greater than one. Yellow dashed lines mark IM components with SNRs significantly greater than one (n1f1+n2f2; n1=1, n2=+-1,+-2,+-3,+-4 as well as n1=2, n2=-1,+2) and black dashed lines mark IM components with SNRs not significantly greater than one.

After establishing that both tagging frequencies and their IM components are present in the data, we examined their spatial distribution on the scalp, averaged across all trials. We expected to find strongest SSVEP amplitudes over the occipital region (as the primary visual cortex is known to be a principal source of SSVEP (32)) and strongest SWIFT amplitudes over more temporal and parietal regions (as they have been shown to increasingly activate higher areas in the visual pathway (21). IM components, in contrast, should originate from local processing units which processes both SSVEP and SWIFT inputs. Under the predictive coding framework, predictions are projected to lower levels in the cortical hierarchy where they are integrated with sensory input. We therefore reasoned that IM signals will be found primarily over occipital regions.

SSVEP amplitude SNRs were strongest, as expected, over the occipital region (Fig. 3A). For SWIFT, highest SNRs were found over more temporo- and centro-parietal electrodes (Fig. 3B). Strongest SNR values for the IM components were indeed found over occipital electrodes (Fig. 3C). To better quantify the similarity between the scalp distributions of SSVEP, SWIFT and IM frequencies we examined the correlations between the SNR values across all channels (n=64). We then examined whether the correlation coefficients for the comparison between the IMs and the SSVEP were higher than the correlation coefficients for the comparison between the IMs and the SWIFT. To do so, we applied the Fisher’s r to z transformation and performed a Z-test for the difference between correlations. We found that the distributions of all IM components were significantly more correlated with the SSVEP than with the SWIFT distribution (z= 6.44, z=5.52, z=6.5 and z= 6.03 for f1+f2, f1-f2, f1+2f2 and f1-2f2, respectively; two-tailed, FDR adjusted p < 0.01 for all comparisons; Fig. S1).

**Figure 3.**
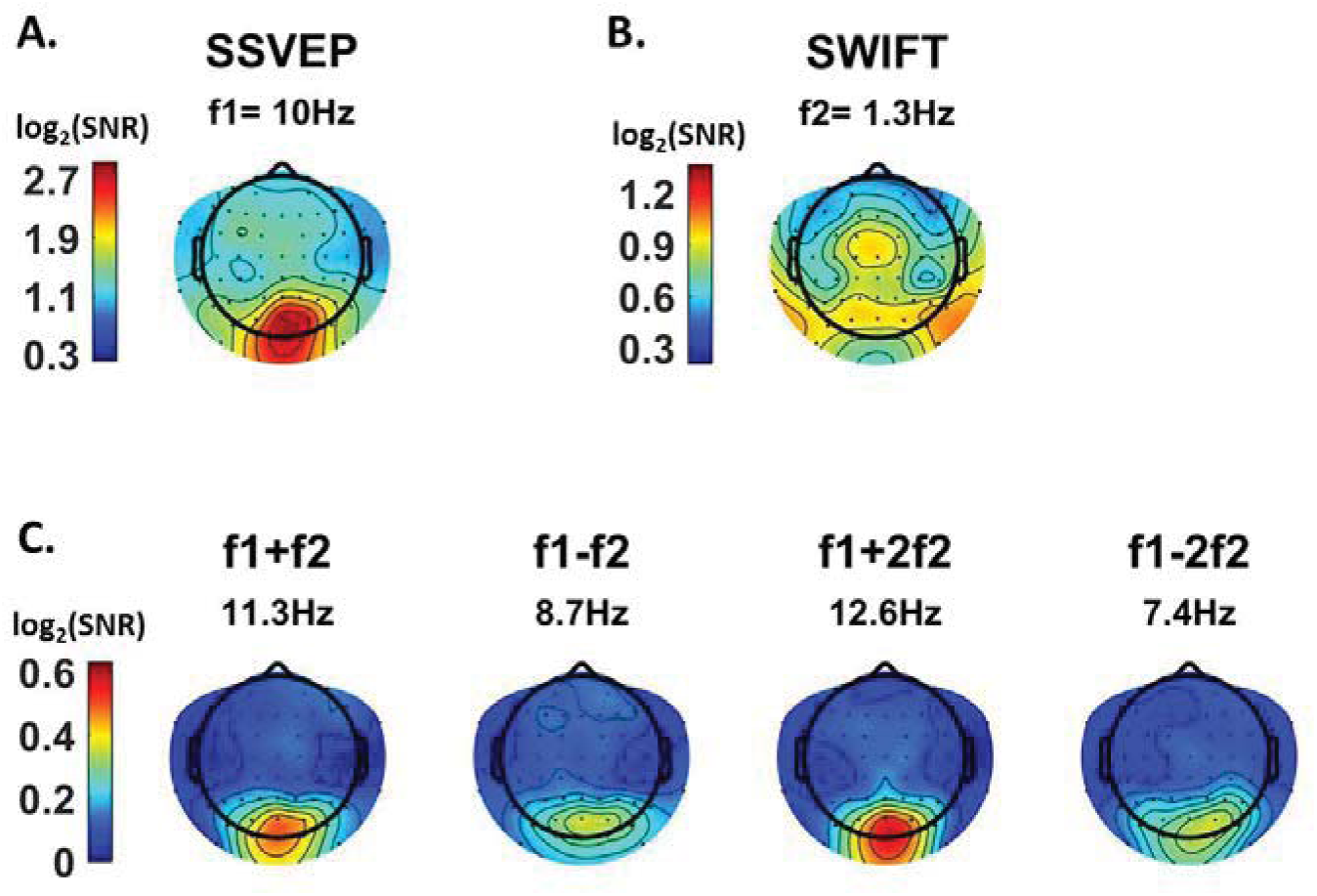
Scalp distributions Topography maps (log2(SNR)) for SSVEP (f1=10Hz) (A), SWIFT (f2= 1.3Hz) (B), and four IM components (f1+f2, f1-f2, f1+2f2 and f1-2f1) (C). SSVEP SNRs were generally stronger than SWIFT SNRs, which in turn were stronger than the IM SNRs (note the different colorbar scales).

As further detailed in the Discussion, we suggest that this result is consistent with the notion that top-down signals (as tagged with SWIFT) are projected to occipital areas, where they are integrated with SSVEP-tagged signals.

The final stage of our analysis was to examine the effect of certainty on the SSVEP, SWIFT and IM signals. Expectations and certainty are suggested to play a significant role in perceptual inference as they shape top-down signals. If the IM components observed in our data reflect a perceptual process in which bottom-up sensory evidence are integrated with top-down predictions, we should expect them to be modulated by the level of certainty about the upcoming stimuli (here, whether the next stimulus would be a face or house image). To test this hypothesis we modulated certainty levels across trials by varying the proportion of house and face images presented.

Using likelihood ratio tests with linear mixed models (see Methods) we found that certainty levels indeed had a different effect on the SSVEP, SWIFT and IM signals (Fig. 4 and Table 1).

**Figure 4.**
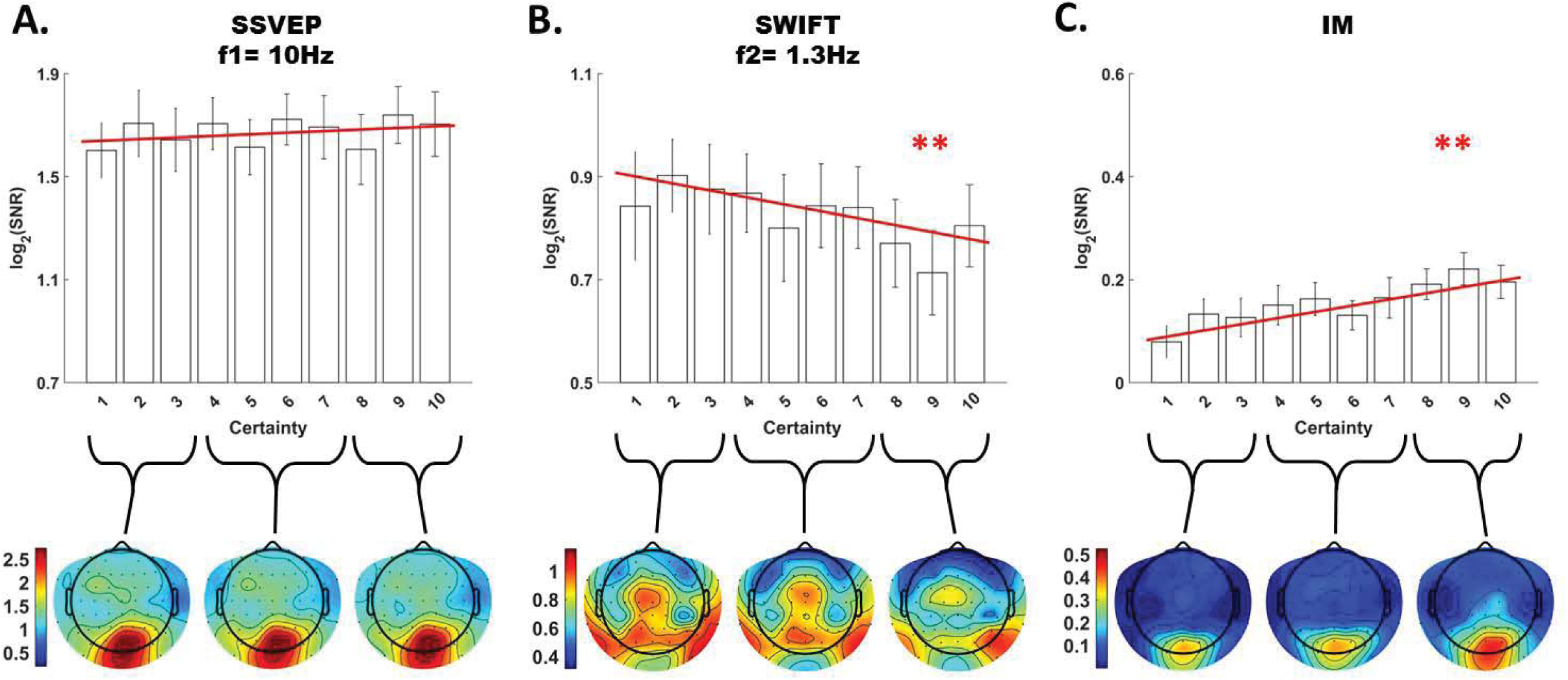
Modulation by certainty Bar plots of signal strength (log of SNR, averaged across 30 posterior channels and 17 participants) as a function of certainty levels for SSVEP (A), SWIFT (B) and IMs (averaged across the 4 IM components) (C). Red lines show the linear regressions for each frequency category. Slopes that are significantly different from 0 are marked with red asterisks (** for p<0.001). While no significant main effect of certainty was found for the SSVEP (p > 0.05), a significant negative slope for was found for the SWIFT, and a significant positive slope was found for the IM. Error bars are SEM across participants. Bottom) Topo-plots, averaged across participants, for low certainty (averaged across bins 1-3), medium certainty (averaged across bins 4-7) and high certainty (averaged across bins 8-10) are shown for SSVEP (A), SWIFT (B) and IM (averaged across the 4 IM components) (C).

**Table 1.**
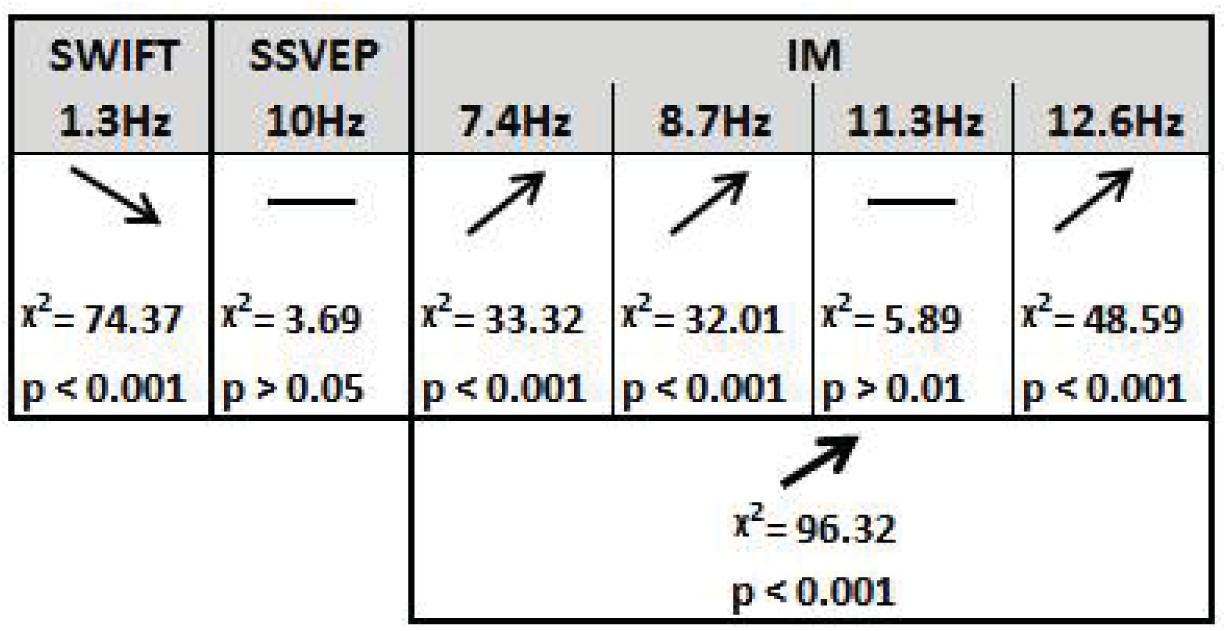
Summary of the linear mixed-effects (LME) modelling. We used LME to examine the significance of the effect of certainty for SSVEP (f1= 10Hz), SWIFT (f2=1.3Hz) and IM (separately for f1-2f2, f1-f2, f1+f2, and f1+2f2, as well as across all 4 components) recorded from posterior ROI electrodes. The table lists the direction of the effects, x^2 value and FDR-corrected p-value from the likelihood ratio tests (See Methods).

First, SSVEP (log of SNR at f1=10Hz) was not significantly modulated by certainty (all Chi square and p-values are shown in Table 1). This result is consistent with the interpretation of SSVEP as mainly reflecting low-level visual processing which should be mostly unaffected by the degree of certainty of the incoming signals.

Second, the SWIFT signals (log of SNR at f2=1.3Hz) significantly decreased in trials with higher certainty. This is consistent with an interpretation of SWIFT as being related to the origin of top-down signals which are modulated by uncertainty. Specifically, better predictions would elicit less prediction error.

Critically, the IM signals were found to increase as a function of increasing certainty for three of the four IM components (f1-2f2=7.4Hz, f1-f2=8.7Hz, and f1+2f2=12.6Hz though not for f1+f2=11.3Hz; Table 1). The effect remained highly significant also when including all four IM components in one model. Indeed, this is the effect we would expect to find if IMs reflect the efficacy of integration between top-down, prediction-driven signals and bottom up sensory input: in ‘high-certainty’ trials the same image appeared in the majority of cycles, allowing for the best overall correspondence between predictions and sensory evidence.

In addition, we found significant interactions between the level of certainty and the different frequency categories (SSVEP/SWIFT/IM). The certainty slope was significantly higher for the IM than for SSVEP (χ^2^ = 12.49, *p* < 0.001) and significantly lower for SWIFT than for SSVEP (χ^2^ = 64.45, *p* < 0.001).

## 3. DISCUSSION

Key to perception is the ability to integrate neural information derived from different levels of the cortical hierarchy (33, 34). The goal of this study was to identify neural markers for the integration between top-down and bottom-up signals in perceptual inference and to examine how this process is modulated by certainty. Hierarchical Frequency Tagging combines the SSVEP and SWIFT methods that have been shown to predominantly tag low levels (V1/V2) and higher, semantically rich levels in the visual hierarchy, respectively. We hypothesised that these signals reflect sensory evidence (or prediction errors) and top-down predictions. Critically, we measured intermodulation (IM) components as an indicator of integration between these signals and hypothesised that they reflect the level of integration between top-down predictions (of different strengths) and bottom-up sensory evidence.

We found significant frequency-tagging for both the SSVEP and SWIFT signals, as well as at various IM components (Fig. 2). This confirms our ability to simultaneously use two tagging methods in a single paradigm and, more importantly, provides evidence for the cortical integration of the SWIFT- and SSVEP-tagged signals. Indeed, the scalp topography for the three frequency categories (SSVEP, SWIFT and IMs) were, as we discuss further below, largely consistent with our hypotheses (Fig. 3) and importantly, they all differed in the manner by which they were modulated by the level of certainty regarding upcoming stimuli. While SSVEP signals were not significantly modulated by certainty, the SWIFT signals decreased and the IM signals increased as a function of increasing certainty (Fig. 4). In the following discussion we examine how these results support the predictive coding framework.

The notion of perceptual inference and the focus on prior expectations goes back as far as Ibn al Haytham in the 11th century who noted that “Many visible properties are perceived by judgment and inference in addition to sensing the object’s form” (35). Contemporary accounts of perception treat these ideas in terms of Bayesian inference and predictive coding (1–4).

Under the predictive coding framework, hypotheses about the state of the external world are formed on the basis of prior experience. Predictions are generated from these hypotheses, which are then projected to lower levels in the cortical hierarchy, and continually tested and adjusted in light of the incoming, stimulus-driven, information. Indeed, the role of top-down signals in perception has been demonstrated in both animal and human studies (9, 13). The elements of the sensory input that cannot be explained away by the current top-down predictions are referred to as the prediction error (PE). This PE is suggested to be the bottom-up signal that propagates from lower to higher levels in the cortical hierarchy until it can be explained away, allowing for subsequent revisions of higher-level parts of the overall hypotheses. Indeed, the notion of PEs has been validated by numerous studies (31, 36-39) and several studies suggest that top-down and bottom up signals can be differentiated in terms of their typical oscillatory frequency bands (15, 18, 40). Perception, under the predictive coding framework, is achieved by an iterative process that singles out the hypothesis that best minimizes the overall prediction error across multiple levels of the cortical hierarchy while taking prior learning, the wider context, and precision estimations into account (2). Constant integration of bottom-up and top-down neural information is therefore understood to be a crucial element in perception (1, 33, 34).

The SSVEP method predominantly tags activity in low levels of the visual hierarchy and indeed highest SSVEP SNRs were measured in our design over occipital electrodes (Fig. 3). We showed that the SSVEP signal was not significantly modulated by certainty (Fig. 4A). These findings suggest that the SSVEP reflects bottom-up sensory evidence, which is consistently tagged throughout the experiment and should not strongly depend on top-down predictions.

The SWIFT method, in contrast, has been shown to increasingly tag higher areas along the visual pathway which process semantic information (21), and we indeed found highest SNRs with the SWIFT in this experiment to distribute over more temporal and parietal electrodes (Fig. 3). Since the activation of these areas depends on image recognition (20), we hypothesised that contrary to the SSVEP, the SWIFT signal should show greater dependency on top-down expectations and certainty (Fig. 4B). Because identifiable images appeared in our stimuli at the SWIFT frequency (1.3Hz), it is reasonable to assume that both predictions and prediction errors would be elicited and tagged at this frequency. With our current paradigm we therefore cannot differentiate between the tagging of prediction and prediction error signals. Nevertheless, various studies have previously demonstrated that highly predictable stimuli tend to evoke reduced neural responses (31, 41, 42). Since PEs reflect the elements of sensory evidence that cannot be explained by predictions, such reduced neural responses have been suggested to reflect decreased PE signals (31). In line with these studies, we suggest that SWIFT SNR decreases as certainty levels increase (Fig. 4B) because SWIFT-tagged signals are mainly weighted by PE signals.

The intermodulation (IM) marker was employed because studying perception requires not only distinguishing between top-down and bottom-up signals but also examining the integration between such signals: in predictive coding terms, the integration between ‘state-units’ and ‘error-units’ within and between different levels of the cortical hierarchy (2). Accordingly, the strength of the Hierarchical Frequency Tagging (HFT) paradigm is in its potential ability to obtain, through the occurrence of IM, a direct electrophysiological measure of integration between signals derived from different levels in the cortical hierarchy.

The scalp distributions of the IM components were more strongly correlated to the distribution of the SSVEP (f1= 10Hz) rather than to the SWIFT (f2= 1.3Hz) (Fig. 4). This pattern supports the notion that the IM components in our Hierarchical Frequency Tagging (HFT) paradigm data reflect the integration of signals generated in SWIFT-tagged areas which project to, and are integrated with, signals generated at lower levels of the visual cortex, as tagged by the SSVEP. This is consistent with the predictive coding framework in which predictions generated at higher levels in the cortical hierarchy propagate to lower areas in the hierarchy where they can be tested in light of incoming sensory-driven evidence.

Importantly, in terms of the dependence on levels of certainty or predictability, we showed that, contrary to SWIFT SNR, IM SNRs *increased* as a function of certainty. We suggest that this result once again lends support specifically to the predictive coding framework: Testing a hypothesis requires the probabilistically sensitive comparison between its associated prediction and the sensory-driven evidence. Higher levels of certainty in our stimuli were associated with greater predictability of upcoming images and a greater overall match throughout the trial between predictions and sensory evidence. The increase in IM SNRs in our data may therefore reflect the efficient integration of predictions and sensory evidence that should be expected when much of the upcoming stimuli is highly predictable. In contrast, when uncertainty increases, the integration diminishes since when the stimuli are less predictable the sensory evidence is harder to explain away.

Several studies have demonstrated a relationship between IM components and perception (27–29). In all of these studies, the reported increase in IM signal strength potentially reflects the integration of different input elements within a neural representation. However, the strength of Hierarchical Frequency Tagging is in its ability to simultaneously tag both bottom-up *and* top-down inputs to the lower visual areas. This is important because if the ‘hypothesis-testing’ stage is to be thought of as a function, then the ‘prediction’ and the ‘evidence’ would be the top-down and bottom-up inputs, respectively, while the PE would be the output (which in turn serves as the evidence for the higher level in the hierarchy). The IM signals, in our paradigm, would then reflect the crux of the hypothesis-testing function, namely, the comparison of prediction and sensory signals, or the integration between state-units and error-units.

Overall, the evidence we have presented plausibly demonstrates our ability, using the novel HFT technique, to obtain a direct physiological measure of the integration of information derived from different levels of the cortical hierarchy during perception. Supporting the predictive coding account of perception, our results suggest that top-down, semantically tagged signals are integrated with bottom-up sensory-driven signals, and this integration is modulated by the level of uncertainty about the causes of the perceived input.

## 4. METHODS

### 4.1 Stimulus construction

#### 4.1.1 SSVEP and SWIFT

In steady-state-visual-evoked-potentials (SSVEP) studies, the intensity (luminance or contrast) of a stimulus is typically modulated over time at a given frequency, *F* Hz (i.e. the ‘tagging frequency’). Peaks at the tagging-frequency, *f* Hz, in the spectrum of the recorded signal are thus understood to reflect stimulus-driven neural activity. However, the use of SSVEP methods impose certain limitations for studying perceptual hierarchies. When the contrast or luminance of a stimulus is modulated over time, then all levels of the visual hierarchy are entrained at the tagging frequency. Thus, it becomes difficult to dissociate frequency tagging related to low-level feature processing from that related to high-level semantic representations.

Semantic wavelet-induced frequency-tagging (SWIFT) overcomes this obstacle by scrambling image sequences in a way that maintains low-level physical features while modulating mid to high-level image properties. In this manner, SWIFT has been shown to constantly activate early visual areas constant while selectively tagging high-level object representations both in EEG (20) and fMRI (21).

The method for creating the SWIFT sequences is described in detail elsewhere (20). In brief, sequences were created by cyclic wavelet scrambling in the wavelets 3D space, allowing to scramble contours while conserving local low-level attributes such as luminance, contrast and spatial frequency. First, wavelet transforms were applied based on the discrete Meyer wavelet and 6 decomposition levels. At each location and scale, the local contour is represented by a 3D vector. Vectors pointing at different directions but of the same length as the original vector represent differently oriented versions of the same local image contour. Two such additional vectors were randomly selected in order to define a circular path (maintaining vector length along the path). The cyclic wavelet-scrambling was then performed by rotating each original vector along the circular path. The inverse wavelet transform was then used to obtain the image sequences in the pixel domain. By construction, the original unscrambled image appeared once in each cycle (1.3Hz). The original image was identifiable briefly around the peak of the embedded image, as has been demonstrated psychophysically (21).

#### 4.1.2 SWIFT-SSVEP trial

SWIFT sequences were created from a pool of grayscale images of houses and faces (28 each, downloaded from the Internet using Google Images (https://www.google.com/imghp) to find images with “free to use, share or modify, even commercially” usage rights; Fig. 1A-B). Each trial was constructed using one house and one face sequence, randomly selected from the pool of sequences (independently from the other trials administered to the participant). In order to preserve low level image attributes within a given trial, regardless of the specific image presented in each cycle, we created and merged additional ‘noise’ sequences in the following way. First, we randomly selected one of the scrambled frames from each of the original SWIFT sequences. Then, we created noise sequences by applying the SWIFT method on each of the selected scrambled frames. In this way, each original ‘image’ sequence had a matching ‘noise’ sequence. Finally, within each trial, each ‘image’ sequence was alpha blended with the ‘noise’ sequences of the other image with equal weights in luminance value at each pixel at any given frame (Figure 1C, ‘image’ sequences are surrounded by solid squares and ‘noise’ sequences with dashed squares). This way, for example, cycles in which a face image was to appear contained the face image superimposed with a house noise sequence (Figure 1C, right side).

This way, the overall low level visual attributes were constant across all frames in the trial regardless of the identifiable image in each cycle, ensuring that the frequency tagged signal would not reflect local physical input properties.

To construct each trial, 65 cycles (each being of either the face or the house image cycle) were presented repeatedly. The resulting trial was a 50 second ‘movie’ in which images peaked in a cyclic manner (F2=1.3Hz) (Figure 1D) in a pseudorandom order. A global sinusoidal contrast modulation at F1=10Hz was applied on the whole movie sequence to evoke the SSVEP.

### 4.2 Participants and Procedure

A total of 27 participants were tested for this study (12 females; mean age = 28.9 y, std = 6.6). Participants gave their written consent to participate in the experiment. Experimental procedures were approved by the Monash University Human Research Ethics Committee.

Participants were comfortably seated with their head supported by a chin rest 50cm from the screen (CRT, 120HZ refresh rate) in a dimly lit room. Sequences were presented at the center of the screen over a grey background and participants were asked to keep their fixation at the center of the display. Participants were asked to minimise blinking or moving during each trial, but were encouraged to do so if needed in the breaks between each 50-sec trial.

A total of 56 such 50-sec trials were presented to each participant. Importantly, the proportion of house and face images varied over trials, spanning the full possible range (pseudorandomly selected such that a particular proportion was not repeated within each participant). Each trial therefore varied in the level of certainty associated with upcoming images.

In order to verify that the participants engaged with the task, a sentence appeared on the screen before each trial instructing them to count either the number of house or face presentations. Trials began when the participant pressed the spacebar. They used the keyboard at the end of each trial to enter the number of images counted. These responses were recorded and used later to exclude poorly-performing participants from the analysis. A 2-3 minute rest break was introduced after every 14 trials. Continuous EEG was acquired from 64 scalp electrodes using a Brain Products BrainAmp DC system. Data were sampled at 1000 Hz for 23 participants and at 500 Hz for the remaining 4 participants.

### 4.3 Data analysis

Data processing was performed using the EEGLAB toolbox (43) in MATLAB. All data sampled at 1000Hz were resampled to 500Hz. A high-pass filter was applied at 0.6Hz.

#### 4.3.1 Exclusion criteria

We defined two criteria to exclude participants from the analysis. First, we excluded participants who had poor counting accuracy because we cannot be sure if these participants were attentive throughout the task. For this purpose, we calculated correlations for each participant between their responses (number of image presentations counted in each trial) and the actual number of cycles in which the relevant image was presented. We excluded five participants whose correlation value r was lower than 0.9 (Fig. 6A).

**Figure 6.**
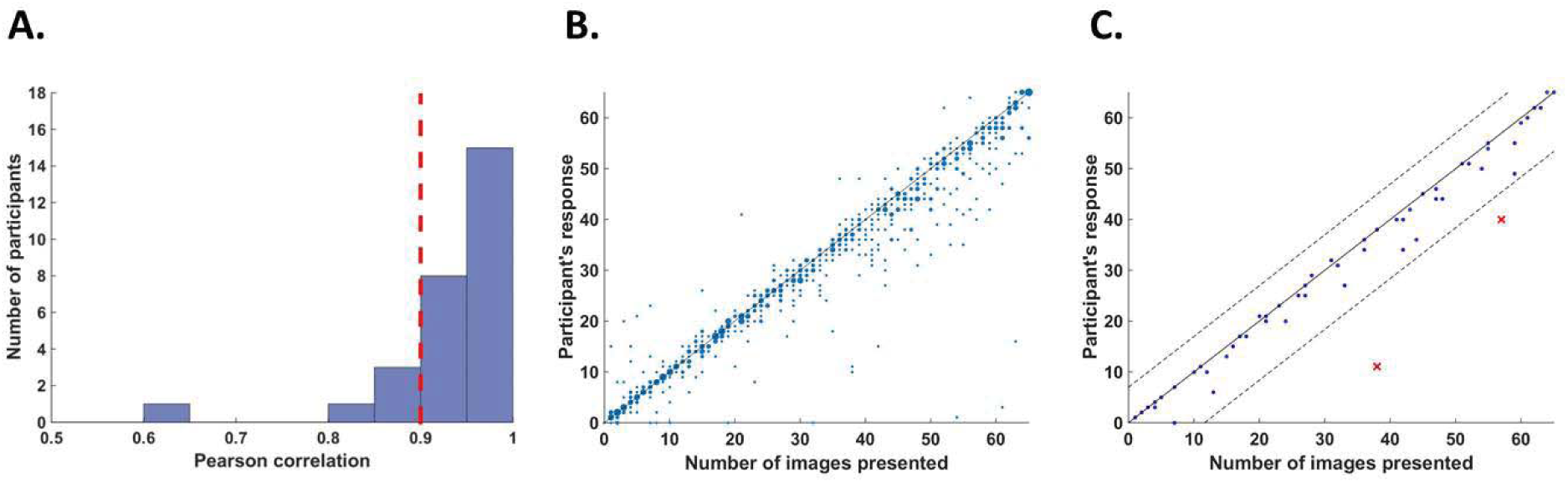
Behavioral performance. (A) Histogram across all participants for counting accuracy measured as the correlation between the participant’s response (number of image presentations counted in each trial) and the actual number of presentations. Five participants with a counting accuracy below r= 0.9 (a vertical red dashed line) were excluded from the analysis. (B) Scatter plot showing responses across 56 trials for all participants included in the analysis. The size of each dot corresponds to the number of occurrences at that point. (C) An example scatter plot for a single participant demonstrating the within-participant exclusion criterion for single trials. The solid line (y=x) illustrates the theoretical location of accurate responses. For each trial, we calculated the distance between the participant’s response and the actual number of cycles in which the relevant image was presented (i.e., the distance between each dot in the plot and the solid line). The within-participant cutoff was then defined as ±2.5 standard deviations from the mean of this distance. Dashed lines mark the within-participant cutoff for exclusion of single trials.

The second criterion was based on the quality of EEG recordings. Sample points were regarded as being noisy if they were either greater than ±80µV, contained a sudden fluctuation greater than 40µV from the previous sample point, or if the signal was more than ±6 std from the mean of the trial data in each channel. Cycles in which over 2% of sample points were noisy were regarded as noisy cycles. For each channel, all sample points within the noisy cycles were replaced by the mean signal across the trial. Participants for which over 10% of cycles were noisy were excluded from the analysis. Five additional participants were excluded on the basis of this criterion for poor EEG recording (on average, 37% of cycles were noisy for these participants). A total of 17 remaining participants were included in the analysis.

In addition, we excluded within-participant subsets of trials. For each participant, we calculated the mean and standard deviation of the difference between the participant’s response (count) and the number of cycles in which the relevant image was presented. We then excluded all trials in which the participant’s response fell further than 2.5 standard deviations from his mean accuracy (e.g., Fig. 2C). From this criterion, we excluded 5.5% of the trials (52 out of 952 trials in total, 0-5 trials out of 56 for any individual participant).

#### 4.3.2 Spectral analysis

EEG signal amplitude was extracted at the tagging and intermodulation frequencies by applying the Fourier transform (FFT) over each trial (50s, 25,000 sample-points, frequency resolution = 0.02 Hz). Signal-to-noise ratios (SNR) at frequency f was computed by dividing the amplitude at f by the mean amplitude across 20 neighbouring frequencies (from f-0.2Hz to f-0.02Hz and from f+0.02Hz to f+0.2Hz) (34, 44).

##### 4.3.2.2 Intermodulation components

IM components include all linear combinations of the fundamental frequencies that comprise the input signal (n1f1 + n2f2, n=±1,±2,±3…). While a large number of potential IM components exist in our data, we focused our analysis on the four lowest-order components (f1-2f2=7.4Hz, f1-f2=8.7Hz, f1+f2=11.3Hz and f1+2f2=12.6Hz, where f1=10Hz and f2=1.3Hz).

#### 4.3.3 Statistical analysis

For analysis of the modulatory effects of certainty we used RStudio (RStudio Team (2015). RStudio: Integrated Development for R. RStudio, Inc., Boston, MA. http://www.rstudio.com/). and lme4 (45) to perform linear mixed-effect analysis of the data. Eight frequencies of interest were analysed: f2=1.3Hz and 2f2=2.6Hz (SWIFT and harmonic), f1=10Hz and 2f1=20Hz (SSVEP and harmonic), and f1-2f2=7.4Hz, f1-f2=8.7Hz, f1+f2=11.3Hz and f1+2f2=12.6Hz (IM components). We used log_2_(amplitude SNR) as the dependant variable for all analyses. We chose this transformation because the amplitude SNR has a lower bound of 0 and does not distribute normally. The distribution of log_2_(SNR) on the other hand is closer to a normal distribution and allows for better homoscedasticity in the linear models.

In order to examine the modulatory effect of certainty, we divided trials into 10 certainty bins ranging from 1 (lowest certainty) to 10 (highest certainty). Bin limits were defined in terms of the percentage of cycles at which the more frequent image appeared, thus creating 5%-wide bins (trials in which the frequent image appeared in 50-55%, 55-60%, … and 95-100% of cycles are defined as bin 1, 2, … and 10, respectively).

Different statistical models were applied for each of the three levels of analysis performed: 1) within each of 6 frequencies of interest (e.g., f1, f2, f1+f2, etc.), 2) within the IM category (f1-2f2, f1-f2, f1+f2 and f1+2f2) and 3) between frequency categories (SSVEP/SWIFT/IM). All analyses were performed on a posterior ROI (30 electrodes) including all centro-parietal (CPz and CP1-CP6), temporo-parietal (TP7-TP10), parietal (Pz and P1-P8), parieto-occipital (POz, PO3-PO4, and PO7-PO10) and occipital (Oz,O1 and O2) electrodes. Channels were added to all models as a random effect. All random effects allowed for both random intercepts and slopes.

To examine if certainty had a significant modulatory effect within each frequency of interest, the first level of analysis included certainty as the fixed effect, and channel nested within participants as the random effect. To examine if there was a main effect for certainty within each frequency category (SSVEP/SWIFT/IM), the second level of analysis included certainty as the fixed effect, and frequency nested within channel nested within participants as the random effect. To examine if the main effect of certainty differed between frequency categories (i.e. a significant interaction between certainty and frequency category), the third level of analysis included certainty, frequency category and a certainty-category interaction as the fixed effects, and frequency nested within frequency category nested within channel nested within participants as the random effect.

To test for the significance of a given factor or interaction, we performed likelihood ratio tests between the full model, as described above, and the reduced model which did not include the factor or interaction in question (45). When applicable, we adjusted p values using the false discovery rate (46).

## ACKNOWLEDGEMENTS

RK was supported by DP130100194 from the Australian Research Council; NT was supported by FT120100619 and DP130100194 from the Australian Research Council; JH was supported by Australian Research Council grants FT100100322 and DP160102770.

## SUPLEMENTARY INFORMATION

**Figure S1.**
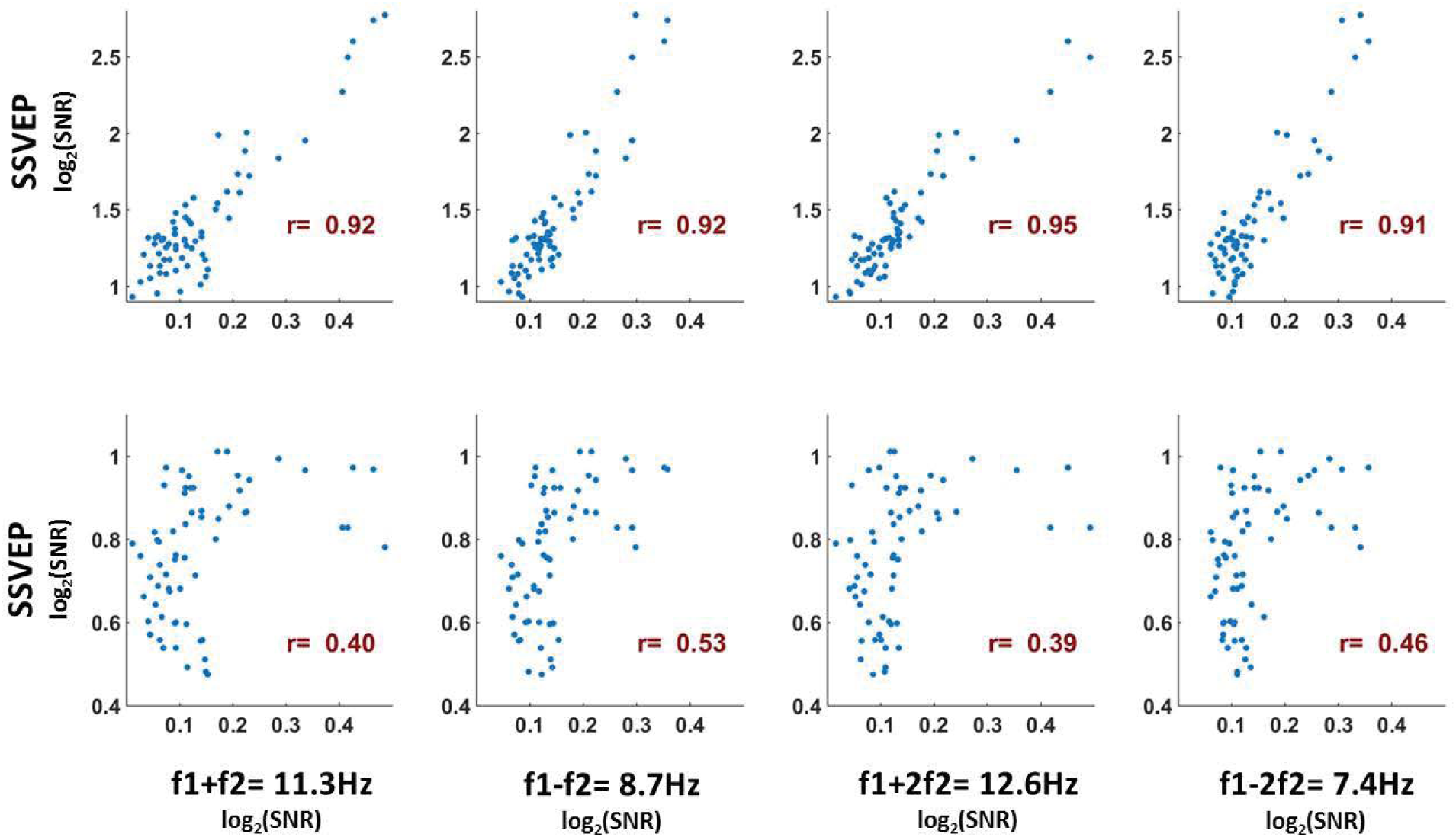
Supplementary Information. As a measure of the similarity between the scalp distributions of the SSVEP, SWIFT and IM frequencies, we examined the Pearson correlation between the mean SNR values across participants for the IM, SSVEP and SWIFT frequencies across all channels (n=64, each point represents the mean SNR for a single channel across 17 participants). To examine whether the correlation coefficients for the comparison between the IMs and the SSVEP were higher than the correlation coefficients for the comparison between the IMs and the SWIFT, we applied the Fisher’s r to z transformation and performed a Z-test for the difference between correlations. We found that the distributions of all IM components were more highly correlated with the SSVEP than with the SWIFT distribution (z= 6.44, z=5.52, z=6.5 and z= 6.03 for f1+f2, f1-f2, f1+2f2 and f1-2f2, respectively; two-tailed, FDR adjusted p < 0.01 for all comparisons).

